# Predicting the Evolution of White Matter Hyperintensities in Brain MRI using Generative Adversarial Networks and Irregularity Map

**DOI:** 10.1101/662692

**Authors:** Muhammad Febrian Rachmadi, Maria del C. Valdés-Hernández, Stephen Makin, Joanna M. Wardlaw, Taku Komura

## Abstract

We propose a Generative Adversarial Network (GAN) model named Disease Evolution Predictor GAN (DEP-GAN) to predict the evolution (i.e., progression and regression) of White Matter Hyperintensities (WMH) in small vessel disease. In this study, the evolution of WMH is represented by the “Disease Evolution Map” (DEM) produced by subtracting irregularity map (IM) images from two time points: baseline and follow-up. DEP-GAN uses two discriminators (critics) to enforce anatomically realistic follow-up image and DEM. To simulate the non-deterministic and unknown parameters involved in WMH evolution, we propose modulating an array of random noises to the DEP-GAN’s generator which forces the model to imitate a wider spectrum of alternatives in the results. Our study shows that the use of two critics and random noises modulation in the proposed DEP-GAN improves its performance predicting the evolution of WMH in small vessel disease. DEP-GAN is able to estimate WMH volume in the follow-up year with mean (std) estimation error of −1.91 (12.12) *ml* and predict WMH evolution with mean rate of 72.01% accuracy (i.e., 88.69% and 23.92% better than Wasserstein GAN).

## 1 Introduction

White Matter Hyperintensities (WMH) are neuroradiological features in T2-weighted and fluid attenuated inversion recovery (T2-FLAIR) brain Magnetic Resonance Images (MRI) that have been associated with stroke and dementia progression [13]. A previous study has shown that the volume of WMH on a patient may decrease (regress), stay the same, or increase (progress) over a period of time [2]. In this study, we refer to these changes as “evolution of WMH”.

Predicting the evolution of WMH is challenging because the rate of WMH evolution varies considerably across studies and patients [10], and factors that influence their evolution are poorly understood [12]. Despite high WMH burden, hypertension, and increasing age have been commonly associated to the evolution of WMH [10], bias in manual delineation of WMH towards progression when the raters are aware of the scans’ time sequence [10] cannot be overlooked.

In this study, we propose an end-to-end training model for predicting the evolution of WMH from *baseline* to the *following year* using generative adversarial network (GAN) [4] and irregularity map (IM) [8,7]. This study differs from other previous studies on predictive modelling focused on the progression of disease and/or its effect (e.g., progression of cognitive decline in Alzheimer’s disease patient [3]). Instead, we are interested in predicting the evolution of specific neuroradiological features in MRI (i.e., WMH in T2-FLAIR). Using our proposed model, clinicians can estimate the size and location of WMH in time to study their progression/regression in relation to clinical health and disease indicators, for ultimately design more effective therapeutic interventions.

The combination of GAN and IM is chosen because of several reasons. We use IM as it enables us to represent the evolution of WMH at voxel level precision using “Disease Evolution Map” (DEM) (full explanation in Section 2). On the other hand, GAN is chosen as it is the *state-of-the-art* method to generate synthetic images. Note that we would like to generate synthetic (fake) DEM which mimics the true (real) DEM. Furthermore, both GAN and IM are unsupervised methods which are not constrained to the availability of labels.

Our main contributions are listed as follows. Firstly, we propose a GAN based model named Disease Evolution Predictor GAN (DEP-GAN) to predict the evolution of WMH. To our best knowledge, this is the first time a GAN based model is proposed for this task. Secondly, we propose the use of two critics which enforce anatomically realistic follow-up images and modifications in DEM. Lastly, we propose modulating an array of random noises to the DEP-GAN to simulate the uncertainty and unknown factors involved in WMH evolution.

## 2 Representation of WMH Evolution using IM

Irregularity map (IM) in our context is a map/image that describes the “irregularity” level for each voxel with respect to the normal brain white matter tissue using real values between 0 and 1 [8]. The IM is advantageous as it retains some of the original MRI textures (e.g., of the T2-FLAIR image intensities), including gradients of WMH. IM is also independent from a human rater or training data, as it is produced using an unsupervised method (i.e., LOTS-IM) [7].

Evolution of WMH at two time points can be obtained by subtracting the baseline IM from the follow-up IM. We call the resulted image “Disease Evolution Map” (DEM) where its values range from −1 to 1. DEM’s values represent the magnitude of the longitudinal change, where negative values mean regression and positive values mean progression. As seen in Fig. 1, DEM calculated from IM represents WMH evolution better than the one calculated from normalised T2-FLAIR MRI. Note how both regression and progression (i.e. dark blue and bright yellow pixels in the figure) are well represented on the DEM from IM at a voxel level precision. T2-FLAIR MRI is not ideal to generate DEM as their voxel values are rather a qualitative representation of the tissue’s spin-spin relaxation time as it decays towards its equilibrium value. Whereas, IM is a quantitative result of assessing how different each voxel is with respect to the ones that make most of the brain tissue voxels (i.e. in T2-FLAIR MRI in this case) [7].

**Fig. 1:**
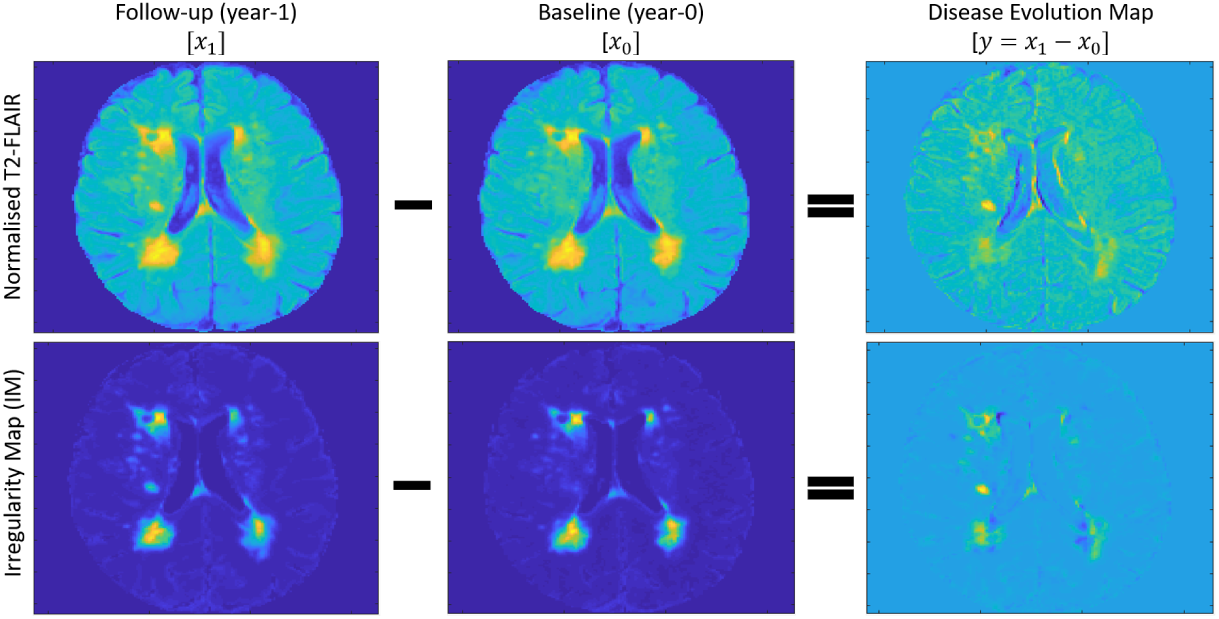
Normalised T2-FLAIR (**top**) and corresponding irregularity maps (IM) (**bottom**) produced by using LOTS-IM [7]. “Disease Evolution Map” (DEM) (**right**) is produced by subtracting baseline images (**middle**) from follow-up images (**left**). In DEM, bright yellow pixels represent positive values (i.e., progression) while dark blue pixels represent negative values (i.e., regression).

### 2.1 MRI data and IM generation

We used MRI data from all stroke patients (*n* = 152) enrolled in a study of stroke mechanisms [12], imaged at three time points (i.e., first time (baseline scan), at approximately 3 months, and a year after). This study uses the baseline and 1-year follow-up MRI data (*s* = *n*×2 = 304), both acquired at a GE 1.5T scanner following the same imaging protocol in [11]. T2-weighted, FLAIR, gradient echo and T1-weighted structural images at all time points were rigidly and linearly aligned using FSL-FLIRT [5]. The resulted working resolution of the images used in this study is 256×256×42 with slice thickness of 0.9375×0.9375×4 mm. The primary study that provided the data used a multispectral method to produce the binary masks used to generate the brain tissue IMs (i.e. intracranial volume (ICV), cerebrospinal fluid (CSF)), and stroke lesions (SL). The image processing protocol is fully explained in [11]. For this study, we obtained IMs from each patient FLAIR imaging data using LOTS-IM [7] with 128 target patches.

## 3 Disease Evolution Predictor GAN (DEP-GAN)

We propose a GAN model for predicting WMH evolution namely disease evolution predictor GAN (DEP-GAN). DEP-GAN is based on the visual attribution GAN (VA-GAN) originally proposed to detect atrophy in T2-W MRI of Alzheimer’s disease [1]. DEP-GAN consists of a generator loosely based on U-Net [9] and two convolutional network critics, where baseline images are feed-forwarded to the generator and fake/real images of follow-up and DEM are feedforwarded to two different critics (see Fig. 2).

**Fig. 2:**
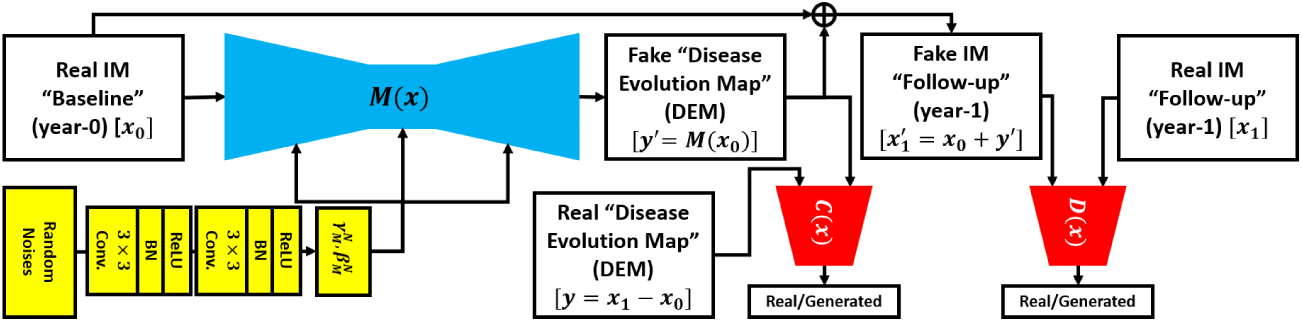
Schematic of the proposed DEP-GAN with 2 discriminators (critics).

Let *x*_0_ be the baseline (year-0) image and *x*_1_ be the follow-up (year-1) image. Then, DEM (*y*) is produced by simple subtraction of *x*_1_ − *x*_0_ = *y*. To generate a fake DEM (*y*′) without *x*_1_, we use a generator function (*M* (*x*)), where *y*′ = *M* (*x*_0_). Thus, a fake follow-up image 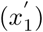 can be easily produced by 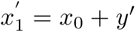. Once *M* (*x*) is well/fully trained, 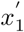 (fake year-1) and *x*_1_ (real year-1) are indistinguishable by a critic function *D*(*x*) while *y*′ (fake DEM) and *y* (real DEM) are also indistinguishable by another critic function *C*(*x*). Full schematic of DEP-GAN’s architecture (i.e., its generator and critics) is provided in the supplementary material.

Unlike VA-GAN model that uses one critic (i.e., *D*(*x*)) [1], two critics (i.e., *D*(*x*) and *C*(*x*)) are used to enforce both anatomically realistic modifications to the follow-up images [1] and anatomical reality of the modifier (i.e., DEM). In other words, we argue that anatomically realistic DEM is also important and essential to produce anatomically realistic (fake) follow-up images.

In this study, we opted for using 2D networks rather than 3D networks because there were only 152 MRI data (subjects) available. For comparison, VA-GAN, which uses 3D networks, used roughly 4,000 MRI data (subjects) for training, yet there was still an evidence of over-fitting [1]. 2D version of VA-GAN itself has been tested on synthetic data [1] and available on GitHub^1^.

### 3.1 Non-deterministic and unknown factors of WMH evolution

The complexity in modelling the evolution of WMH is mainly due to its non-deterministic nature, as the factors that influence it are not fully well known. Previous studies have identified some associated factors (e.g., age, blood pressure, and WMH burden), but their level of influence differs in each study [10,12]. To simulate the non-deterministic and unknown parameters involved in WMH evolution, we propose modulating random noises (*z* ~ 𝒩(0, 1)) to every layer of the DEP-GAN’s generator using Feature-wise Linear Modulation (FiLM) layer [6] (see green block in Fig. 3). In FiLM layer, *γ*_*m*_ and *β*_*m*_ modulate feature maps *F*_*m*_, where subscript *m* refers to *m*^*th*^ feature map, via affine transformation:

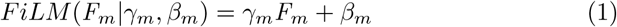

In this study, *γ*_*m*_ and *β*_*m*_ for each residual block (ResBlock) are determined automatically by convolutional layers (yellow blocks in Fig. 3). In this study, the random noises follow Gaussian distribution of *z* ~ 𝒩(0, 1) with the length of 32.

**Fig. 3:**
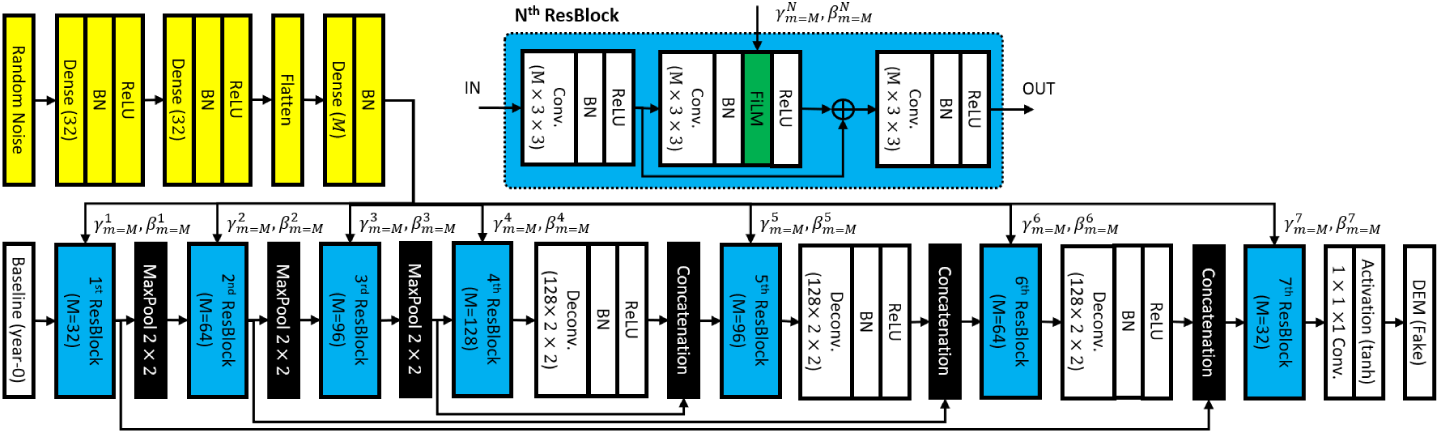
Schematic of DEP-GAN’s generator with Feature-wise Linear Modulation (FiLM) layer [6] (depicted in green block) to simulate the non-deterministic and unknown parameters involved in WMH evolution.

**Fig. 4:**
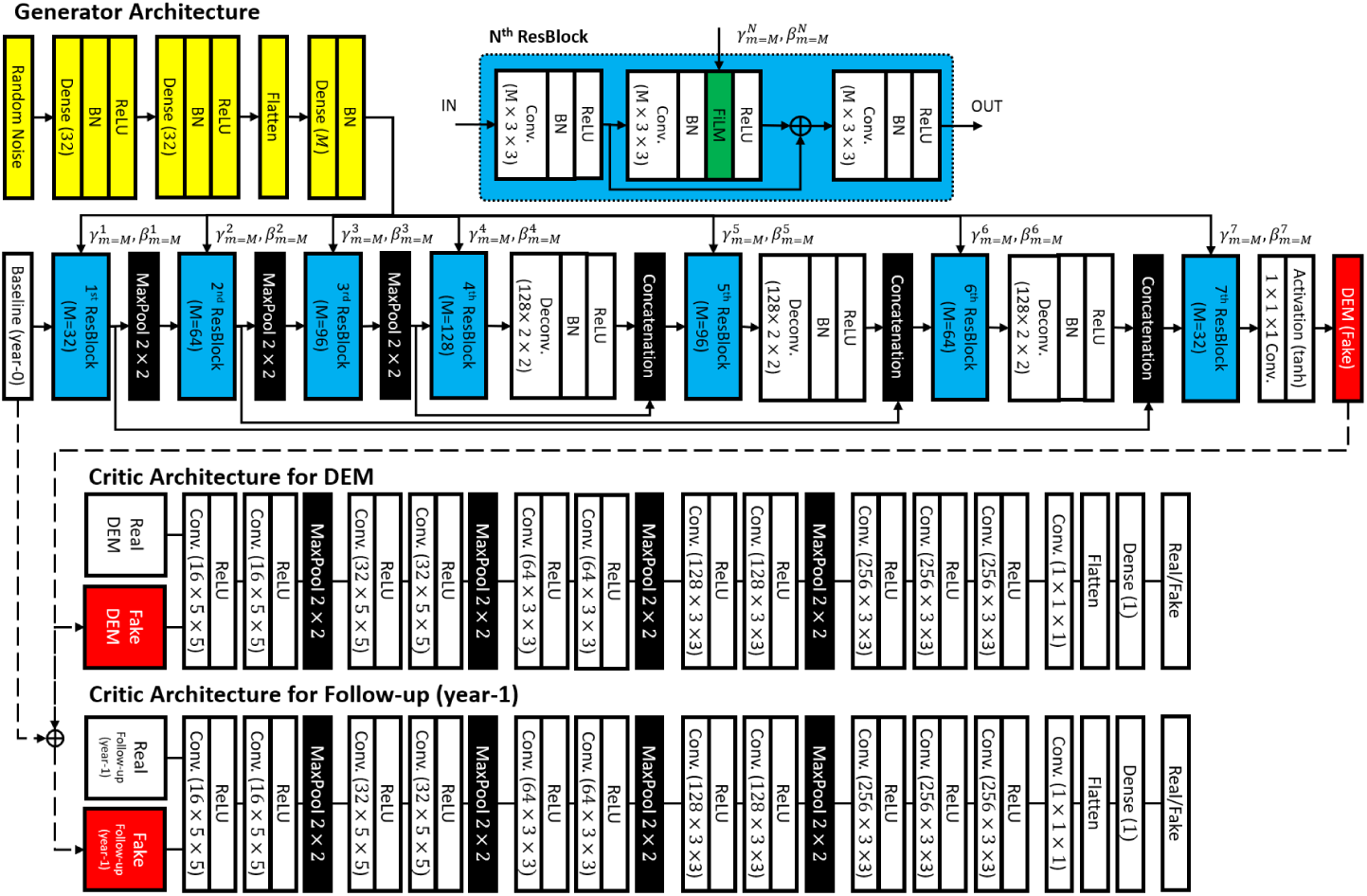
Architecture of generator (**top**) and critics (**bottom**) of DEP-GAN. Note the proposed additional randomness scheme is also depicted where random noises are encoded using convolutional layers (yellow) and then modulated to the generator using FiLM layer (green) inside ResBlock (light blue).

**Fig. 5:**
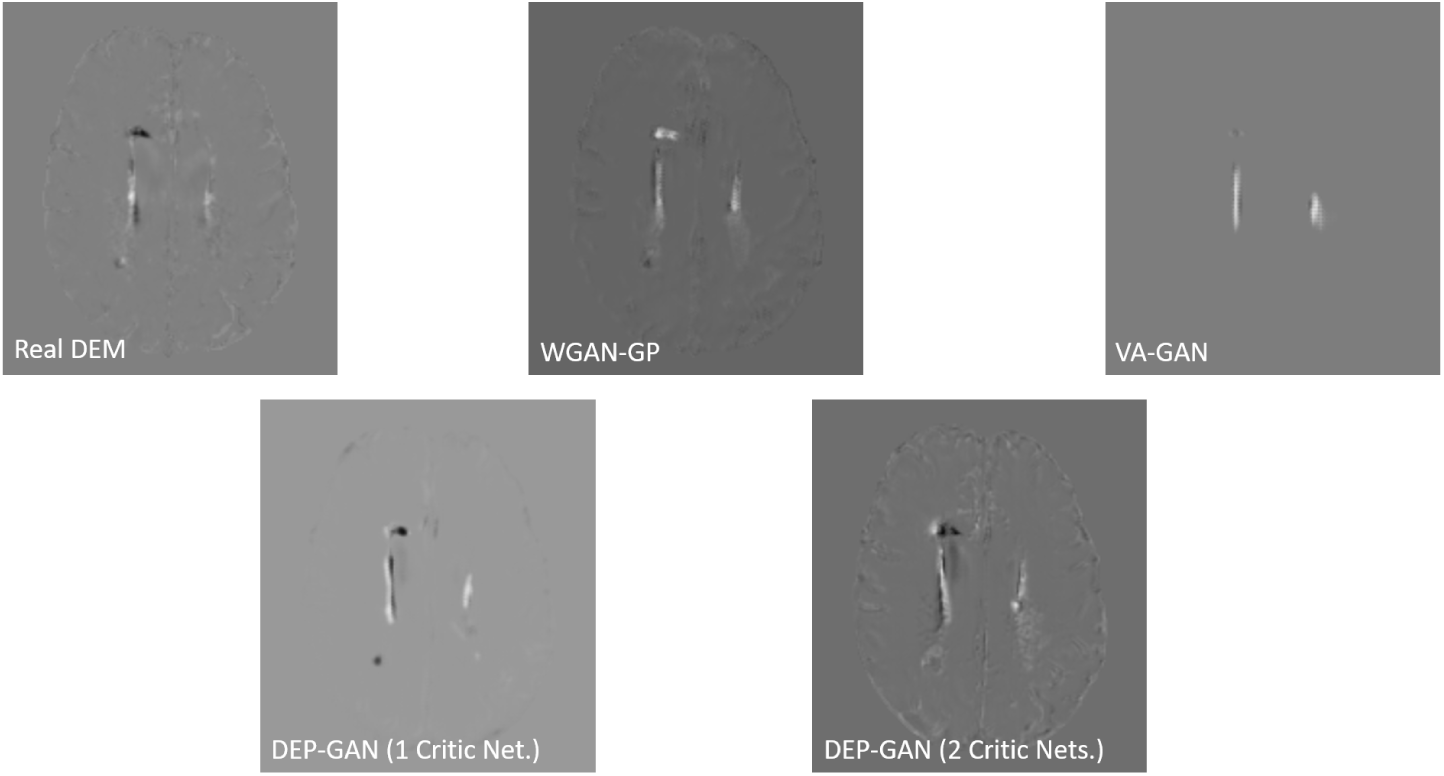
Comparison between real DEM and fake DEM generated from different networks of GANs.

### 3.2 Loss function of DEP-GAN

We build DEP-GAN based on the improved Wasserstein GAN (WGAN-GP) that finds the optimal *M* (*x*) generator function using training approach proposed in [4]. We use a gradient penalty factor of 10 for all experiments. The optimisation of *M* (*x*) is given by the following functions:

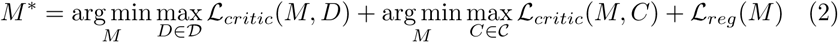

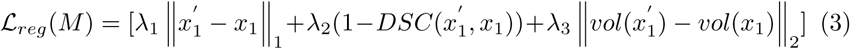

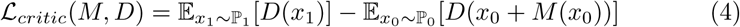

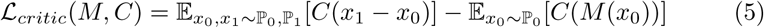

where *x*_0_ baseline images come from underlying distribution of ℙ_0_, *x*_1_ follow-up images come from underlying distribution of ℙ_1_, *M* (*x*_0_) are fake DEM, 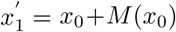 are fake follow-up images, 𝒟 and 𝒞 are the set of 1-Lipschitz functions [1,4], and ‖·‖_1_/ ‖·‖_2_ are the L1/L2 norm.

In summary, to optimise the generator *M* (*x*), we need to optimise Eq. 2, which optimises both critics (i.e., *D*(*x*) and *C*(*x*)) based on WGAN-GP [4], and regularise it with Eq. 3. The regularisation function (Eq. 3) simply says: a) fake follow-up images 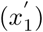 have to be similar to real follow-up images (*x*_1_) based on L1 norm [1], b) WMH segmentation from 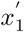 has to be spatially similar to WMH segmentation from *x*_1_ based on the Dice similarity coefficient (DSC), and c) WMH volume (*ml*) from 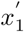 has to be similar to WMH volume from *x*_1_ based on L2 norm. The WMH segmentation of 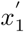 and *x*_1_ is estimated by thresholding their IM values (i.e., irregularity values) to be above 0.178 [7]. Each term in Eq. 3 is weighted by *λ*_1_, *λ*_2_, and *λ*_3_ which equals to 100 [1], 1 and 100 respectively.

## 4 Experiment and Evaluation Setups

We used T2-FLAIR MRI data obtained at two different time points (i.e., 1 year interval) from 152 stroke patients. For testing, we selected 30/152 subject’s data with visible increase (i.e., progression) (19 subjects) and decrease (i.e., regression) (11 subjects) in WMH volume. Thus, data from 122 subjects (i.e., 244 baseline and follow-up MR images) were used for training. As previously described in Section 3, all tested models were trained in 2D (i.e., per slice) manner. Out of all slices from training data, 20% of them were randomly selected for validation. Thus, around 4,000 slices were used in the training process.

From the co-registered scans [11], we generated IM using LOTS-IM [7] with 128 target patches and removed the extracranial tissues and skull from the baseline and follow-up T2-FLAIR images. Then, T2-FLAIR values were normalised between 0 and 1, similar to IM’s values. We also excluded the stroke lesions using the SL masks obtained from previous analyses [2,12] as per [11].

We evaluated WGAN-GP, VA-GAN, and DEP-GAN with 1 critic (for follow-up data only) and compared their performances with DEP-GAN with 2 critics (for follow-up data and DEM). We used our implementation of 2D VA-GAN, following [1]. For WGAN-GP, we modified our implementation of VA-GAN so that its critic learns to distinguish real/fake DEM, not follow-up data. Furthermore, T2-FLAIR was also used as input channel for the generator of DEP-GAN. On the other hand, only IM was feedforwarded to the critics of DEP-GAN/VA-GAN.

Following [1,4], all methods were optimised by updating the parameters of critic(s) and generator in an alternating fashion where (each) critic is updated 5 times per generator update. Furthermore, for the first 25 iterations and every hundredth iteration, critic was updated 100 times per generator update [1]. The generator itself was updated 200 times (i.e., 200 epochs).

In this study, we used 3 evaluation metrics, i.e., 1) estimation error of WMH volume in the follow-up year, 2) prediction error of WMH evolution (i.e., whether WMH in a subject will regress/progress), and 3) Dice similarity coefficient (DSC) (i.e., evaluating the location of WMH evolution). Estimation error of WMH volume (in *ml*) is calculated as 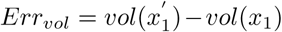. Whereas, to calculate the DSC, we first performed subtraction between WMH segmentation of the baseline image (*x*_0_) from WMH segmentation of the real (*x*_1_) and fake 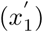 follow-up images. Then, we labelled each voxel as either “Shrink” (Sr.) if it has value below zero, “Grow” (Gr.) if it has value above zero, or “Stay” (St.) if it has value of zero. DSC itself can be computed as *DSC* = ^2*TP*^*/*_(2*TP* +*FP* +*FN*)_ where *TP* is true positive, *FP* is false positive and *FN* is false negative. As previously mentioned, WMH correspond to IM values equal or higher than 0.178 [7] is used as main reference (i.e., ground truth).

## 5 Result

All quantitative evaluations are listed in Table 1, where performance of DEP-GAN-2C (i.e., with 2 critics) is compared to the performance of WGAN-GP, VA-GAN, and DEP-GAN-1C (i.e., with 1 critic). Depiction of real DEM and fake (generated) DEM can be seen in supplementary materials.

**Table 1:** Evaluations of predicting WMH evolution using volumetric error (*Err*_*vol*_), accuracy of evolution prediction (*Pred*_*evo*_), and Dice similarity coefficient (DSC). **Abbreviations**: “std.” for standard deviation, “med.” for median, “Sr.” for shrink, “Gr.” for grow, “St.” for stay, and “Avg.” for average.

| Method | $Err_{vol}$ (ml) | | | | | $Pred_{evo}$ (%) | | | DSC | | | |
| --- | --- | --- | --- | --- | --- | --- | --- | --- | --- | --- | --- | --- |
|  | mean | std. | med. | min | max | Sr. | Gr. | Avg. | Sr. | Gr. | St. | Avg. |
| WGAN-GP [4] | -16.89 | 13.30 | -15.58 | -58.94 | **1.08** | **90.91** | 5.26 | 48.09 | 0.3082 | 0.1600 | 0.6926 | 0.3869 |
| VA-GAN [1] | -18.76 | 12.99 | -16.57 | -59.76 | 1.10 | **90.91** | 0.00 | 45.46 | 0.0409 | **0.2356** | **0.7011** | 0.3259 |
| DEP-GAN-1C | -4.11 | **12.12** | -4.45 | -46.20 | 33.67 | 81.82 | 15.79 | 48.80 | 0.3979 | 0.0862 | 0.6838 | 0.3893 |
| DEP-GAN-2C | **-1.91** | **12.12** | **-1.89** | **-44.35** | 34.73 | 54.55 | **89.47** | **72.01** | **0.4833** | 0.1325 | 0.6606 | **0.4272** |

From Table 1, we can see that DEP-GAN-2C performed better than the other methods in all experiments. Estimating WMH volume, DEP-GAN-2C and DEP-GAN-1C are the best and second-best methods with mean error of estimation −1.91 *ml* and −4.11 *ml* (i.e., 88.69% and 75.67% better than WGAN-GP). Whereas, WGAN-GP and VA-GAN largely underestimated the WMH volume.

DEP-GAN-2C also performed better than the other methods in evolution prediction. DEP-GAN-2C correctly predicted WMH evolution with average rate of 72.01% (i.e., 54.55% for shrink and 89.47% for growth). Whereas, WGAN-GP, VA-GAN, and DEP-GAN-1C failed to predict the growth of WMH most of the time. Note that this correlates with the estimation of WMH volume experiment.

In DSC evaluation, DEP-GAN-2C performed better than the other methods with average DSC of WMH evolution 0.4272 (i.e., 0.4833, 0.1325, and 0.6606 for shrink, growth, and stay respectively). On the other hand, WGAN-GP, VA-GAN, and DEP-GAN-1C performed worse than DEP-GAN-2C where the average DSC were 0.3869, 0.3259, and 0.3893 respectively.

## 6 Discussion and Future Work

In this study, we propose an end-to-end model named DEP-GAN with 2 critics (DEP-GAN-2C) which outperforms WGAN-GP, VA-GAN, and DEP-GAN-1C for predicting the evolution of WMH from 1 time point assessment without any manual WMH label. Based on the results, DEP-GAN-2C had the best performance amongst other methods in estimating both size and location of WMH in the future/follow-up. However, we found that identifying the position of WMH evolution (especially for progression/growth) is the most challenging part of the study as DSC metrics are still low for all methods. From visual inspection of the fake (generated) DEM, we observe that: 1) VA-GAN emphasised major progression/regression, but it neglected minor ones which lead to very low DSC on shrinking WMH; 2) DEP-GAN-1C produced better DEM than VA-GAN thanks to better loss function and simulation of uncertainty/unknown factors, but it does not look realistic like the DEM produced by DEP-GAN-2C; and WGAN-GP produced surprisingly realistic DEM, but it fell short in other evaluations. In the future, the performance of DEP-GAN might be improved by modulating other biomarkers (i.e., non-MRI) to the generator. DEP-GAN code is available at https://github.com/febrianrachmadi/dep-gan-im.

## Acknowledgement

Funds from the Indonesia Endowment Fund for Education (LPDP), Ministry of Finance, Republic of Indonesia (MFR); Row Fogo Charitable Trust (Grant No. BRO-D.FID3668413)(MCVH); Wellcome Trust (patient recruitment, scanning, primary study Ref No. WT088134/Z/09/A); Fondation Leducq (Perivascular Spaces Transatlantic Network of Excellence); EU Horizon 2020 (SVDs@Target); and the MRC UK Dementia Research Institute at the University of Edinburgh (Wardlaw programme) are gratefully acknowledged.

## Footnotes

1 https://github.com/baumgach/vagan-code

## Notes

https://github.com/febrianrachmadi/dep-gan-im

